# *In vitro* reduction of mineral oil hydrocarbons by *Lacticaseibacillus casei* DSM 20011

**DOI:** 10.64898/2026.11.12.766084

**Authors:** Ana Miklavčič Višnjevec, Michela Pellegrini, Chiara Conchione, Laura Barp, Donatella Peressini, Lucilla Iacumin, Sabrina Moret

## Abstract

Mineral oil hydrocarbons (MOH) are food contaminants of increasing concern due to their widespread occurrence in food and potential health risks. Although various physicochemical and biological strategies have been investigated to reduce chemical contaminants in food, the ability of lactic acid bacteria (LAB) to remove MOH has not previously been explored. The present study aimed to evaluate the *in vitro* removal of MOH by *Lacticaseibacillus casei* DSM 20011 under incubation at 37 °C for 4 h with shaking at 100 rpm. Following the screening of different LAB strains, *L. casei* DSM 20011 was selected for *in vitro* evaluation of MOH removal using a representative mineral oil mixture, followed by the determination of MOH by on-line LC–GC-FID. *L. casei* DSM 20011 reduced MOH concentrations by up to approximately 80% (*p* < 0.05), with comparable reductions observed for the MOSH and MOAH fractions and no clear evidence of preferential removal of either fraction. Cell wall adsorption has been proposed as a mechanism contributing to PAH removal by LAB. Given the structural similarity between PAH and the MOAH fraction of MOH, together with the lack of recognized hydrocarbon-degrading enzymes in LAB, adsorption to the bacterial cell wall may also explain the observed MOH removal, although this hypothesis requires experimental confirmation.These findings identify the food-derived *L. casei* DSM 20011 as a promising candidate for the biological reduction of MOH contamination and provide a foundation for future studies investigating the mechanisms underlying MOH removal and validating its effectiveness in complex food matrices.

## 1. Introduction

Mineral oil hydrocarbons (MOH) are contaminants of increasing concern due to their widespread occurrence in food and their potential health risks. MOH comprise a complex mixture of hydrocarbons containing approximately 10 to 50 carbon atoms, and their definition is closely linked to their analytical determination. They originate primarily from petroleum and petroleum-derived products but may also be produced synthetically from coal, natural gas, or biomass. Based on their chemical structure, MOH are classified into two major groups, each comprising a large number of isomers: 1) mineral oil saturated hydrocarbons (MOSH) consisting of linear and branched aliphatic hydrocarbons (alkanes and isoalkanes) together with cyclic saturated hydrocarbons (cycloalkanes), and 2) mineral oil aromatic hydrocarbons (MOAH), comprising hydrocarbons with one or more aromatic rings, predominantly alkylated benzene derivatives (EFSA, 2012; Bratinova *et al*., 2023). MOH can migrate into food from recycled paper and cardboard packaging or be introduced during food processing and through environmental contamination (Bratinova *et al*., 2023; EFSA, 2023; Neukom *et al*., 2008; Van de Ven *et al*., 2018).

Although MOSH are generally considered to pose a relatively low toxicological risk, certain MOAH, particularly compounds containing three or more aromatic rings with a low degree of alkylation, have been associated with genotoxic and carcinogenic effects (EFSA, 2023). Reflecting these health concerns, the European Commission has announced its intention to establish maximum permitted levels of MOAH in food by 2027 (European Commission, 2024).

Various physicochemical and biological strategies have been investigated to reduce chemical contaminants in food. Among biological approaches, lactic acid bacteria (LAB) have attracted considerable attention because of their ability to reduce a wide range of environmental contaminants in food (Abdelshafy et al., 2024). Different studies showed that both viable and inactivated LAB can remove polycyclic aromatic hydrocarbons (PAH), including benzo(a)pyrene (BaP), benzo(a)anthracene (BaA), chrysene (Chr), and benzo(b)fluoranthene (Yousafi et al., 2018; Yousafi et al., 2021; Zhao et al., 2013; Zhang et al., 2023). This removal is believed to occur mainly through physical adsorption onto bacterial cell wall components, particularly peptidoglycans (Zhao et al., 2013). However, whether LAB can also remove MOH has not yet been investigated. Therefore, the present study aimed to evaluate, for the first time, the *in vitro* removal of MOH by *Lacticaseibacillus casei* DSM 20011.

## 2. Experimental

### 2.1 Bacterial strain selection and growth conditions

*Lactiplantibacillus plantarum* DSM 20174, *Lacticaseibacillus paracasei* DSM 20020, *Lacticaseibacillus rhamnosus* DSM 20021, and *Lacticaseibacillus casei* DSM 20011 were initially evaluated with at least two biological replicates.

Based on the results of this initial strain screening, *L. casei* DSM 20011 was selected as the best-performing strain was selected for subsequent experiments. The trials were designed as follows: i) PBS containing the MOH mixture and the bacterial strain to be tested; and two control samples, (ii) PBS containing neither the MOH mixture nor bacteria. The initial strain screening followed the same protocol as that described below for the selected strain, *L. casei* DSM 20011.

The strains, as well as the finally selected, were recovered from stocks stored at −80 °C. Briefly, 100 µL of the frozen stock culture was inoculated into 5 mL of De Man, Rogosa, and Sharpe (MRS) broth (Oxoid, Milan, Italy) and incubated at 30 °C for 24 h. Subsequently, 100 µL of the broth culture was spread onto MRS agar plates and incubated at 30 °C for 48 h under anaerobic conditions. Following confirmation of culture purity, a single colony was inoculated into 4 mL of MRS broth and incubated at 30 °C for 48 h. Then, the bacterial culture was centrifuged at 12,000 × *g* for 10 min. The resulting pellet was washed with 2 mL of sterile physiological saline (9 g NaCl/L), followed by a second centrifugation step at 12,000 × *g* for 10 min. The pellet was then resuspended in 1 mL of sterile physiological saline. The bacterial concentration was estimated based on viable plate counting and by measuring the optical density at 600 nm (OD_600_) using a spectrophotometer (Bio-Rad SmartSpec 3000, Germany). The bacterial suspension (approximately 10 CFU/mL) was used for all subsequent trials.

### 2.2 Preparation of MOH standard working solutions

To evaluate the *in vitro* removal of MOH by LAB, a representative mineral oil mixture (GRAVEX) was used as the model contaminant. GRAVEX consists of a mixture of a naphthenic-based process oil containing approximately 25% of MOAH, supplied by the manufacturer (Udine, Italy). Stock standard solutions was prepared with cyclohexane (10.15 mg/mL). An aliquot of the GRAVEX stock solution was dissolved in 2 mL of 70% ethanol containing 1% Tween® 80 (Sigma-Aldrich, St. Louis, MO, USA) to facilitate dispersion. Subsequently, 200 µL of this solution was added to 4 mL of sterile PBS (Sigma-Aldrich, St. Louis, MO, USA; P4417-100TAB) for the *in vitro* removal assay. In addition, a reference solution of the GRAVEX mixture was prepared in toluene (Merck, Burlington, MA, USA) at the same concentration (47.6 mg/L) used in the *in vitro* assay. This solution was analyzed directly by LC–GC-FID and served as the analytical reference for comparison with samples obtained after the *in vitro* removal assay.

### 2.3 *In vitro* MOH removal assays

The *in vitro* MOH removal assay was performed according to the method described by Yousefi *et al*. (2018), with minor modifications. Briefly, 4 mL of sterile PBS supplemented with 1 mL of the 10 CFU/mL *Lacticaseibacillus casei* DSM 20011 suspension (prepared in physiological saline) and the GRAVEX mineral oil mixture at the concentration 47.6 mg/L. Two control samples were also included: 1) PBS containing the GRAVEX mineral oil mixture but no bacteria and 2) PBS containing neither the GRAVEX mineral oil mixture nor bacteria. All samples were incubated at 37 °C for 4 h with shaking at 100 rpm as was previously described by Yousfi *et al*., 2018, 2021. Following incubation, the samples were transferred to centrifuge tubes previously rinsed with HPLC-grade *n-*hexane and centrifuged at 5,000 rpm for 5 min (Hermle Z 206 A, Wehingen, Germany). The resulting supernatant was collected and subjected to LC–GC-FID analysis for the determination of MOH.

### 2.4 Extraction and LC-GC-FID analysis of MOH

MOH extraction was performed by adding 4 mL of HPLC-grade *n*-hexane (Merck, Burlington, Massachusetts, USA) to the centrifuged samples. The tubes were sealed with aluminum foil, shaken, and sonicated for 5 min. Subsequently, the samples were vortexed for 30 s, followed by a second 5-min sonication step. Then, the samples were centrifuged at 5,000 rpm for 5 min (Hermle centrifuge Z 206 A, Wehingen, Germany). Following phase separation, 100 µL of the clear *n*-hexane layer was transferred to a *n*-hexane-rinsed vial and diluted with 900 µL of HPLC-grade *n*-hexane (1:10, v/v). Finally, 10 µL of the diluted extract was injected into the LC–GC-FID system for MOH analysis.

MOH analysis was performed according to Moret *et al*. (2025) using an online LC–GC-FID system (LC-GC 9000, Brechbühler, Zurich, Switzerland), consisting of a Phoenix 9000 HPLC pump (Brechbühler, Zurich, Switzerland) coupled to a Trace 1310 gas chromatograph (Thermo Fisher Scientific, Waltham, MA, USA). The system was equipped with a PAL LHS2-xt autosampler (CombiPAL, Zwingen, Switzerland) and configured with a dual-channel GC setup, allowing the simultaneous determination of the MOSH and MOAH fractions in a single HPLC run. Chromatographic separation was performed using a LiChrospher Si-60 column (250 × 2.1 mm, 5 μm; DGB, Schlossböckelheim, Germany). The mobile phase initially consisted of 100% *n*-hexane (HPLC grade, distilled before use; Sigma-Aldrich, St. Louis, MO, USA) delivered at a flow rate of 300 μL/min for 0.1 min. A linear gradient was then applied over 0.5 min to a 70:30 (v/v) *n*-hexane/dichloromethane mixture (both HPLC grade, distilled before use; Sigma-Aldrich, St. Louis, MO, USA), while maintaining the flow rate at 300 μL/min. Six minutes after sample injection, the HPLC column was backflushed with dichloromethane at 500 μL/min for 9 min, followed by re-equilibration with *n*-hexane at 700 μL/min for 6.5 min and subsequently at 300 μL/min for 1.5 min. The MOSH and MOAH fractions were automatically transferred into two separate GC columns within the same GC oven via a Y-interface as described by Biedermann and Grob (2009). Briefly, the interface consisted of a glass press-fit connector joining the hydrogen carrier gas line and the LC effluent to a deactivated retention gap. The retention gap was connected through a stainless-steel T-piece to a GC capillary column (10 m × 0.25 mm i.d.) coated with a PS-255 stationary phase (1% vinyl, 99% methylpolysiloxane; film thickness 0.15 μm; Mega, Legnano, Milan, Italy) and to a capillary connected to a solvent vapor exit maintained at 140 °C. Transfer of the LC fractions to the GC was based on the partially concurrent eluent evaporation principle described by Biedermann et al. (2009).

The GC oven temperature was initially maintained at 55 °C for 8.5 min, then increased to 350 °C at a rate of 20 °C/min and held for 5 min. Hydrogen was used as the carrier gas at a constant pressure of 60 kPa, which was increased to 90 kPa during transfer of the LC fractions to the GC. The solvent vapor exit (SVE), kept open during the transfer step, and the FID base block were maintained at 140 °C and 360 °C, respectively. GC performance was verified using an *n*-alkane standard mixture (even-numbered *n*-C10–C40 supplemented with *n*-C50; Restek, Bellefonte, PA, USA). Instrument performance was further monitored using an LC–GC internal standard (IS) solution in toluene (Restek, Bellefonte, PA, USA), containing *n*-C13 (0.15 mg/mL); 1,3,5-tri-*tert*-butylbenzene (TBB), *n*-C11, cyclohexylcyclohexane (CyCy), pentylbenzene (5B), 1-methylnaphthalene (1MN), and 2-methylnaphthalene (2MN) (0.30 mg/mL each); and 5-α-cholestane (Cho) and perylene (Per) (0.60 mg/mL each) as previously reported (Moret *et al*., 2025). Chromatographic data were acquired and processed using Chromeleon™ 7 Chromatography Data System (CDS) software (Thermo Fisher Scientific, Waltham, MA, USA) according to the updated JRC guidance for MOH analysis (Bratinova et al., 2023). Percentage reduction was calculated relative to Control 1 (PBS containing the MOH mixture without LAB).

### 2.5 Statistical analysis

All initial strain screening experiments performed for strain selection were conducted with at least two biological replicates, and the results are presented as the mean ± standard deviation (SD). The main experiment with the selected strain was performed in eight independent replicates (*n* = 8), while each control sample was analyzed in four independent replicates (*n* = 4). Statistical analyses were conducted using STATA/SE 11.0 (StataCorp LP, College Station, TX, USA). As the data were not normally distributed, differences between groups were evaluated using the Wilcoxon–Mann–Whitney test. Statistical significance was set at p < 0.05.

## 3. Results and discussion

### 3.1 Selection of lactic acid bacteria

An initial strain screening of *Lactiplantibacillus plantarum* DSM 20174, *Lacticaseibacillus paracasei* DSM 20020, *Lacticaseibacillus rhamnosus* DSM 20021, and *Lacticaseibacillus casei* DSM 20011 was performed to identify the most promising candidate for *in vitro* MOH removal.. The results of the initial strain screening are presented in **Table S1**.

All four lactic acid bacterial strains reduced the MOSH and MOAH concentrations compared with the control containing the MOH mixture without bacteria, indicating their ability to remove MOH *in vitro*. Among the tested strains, *L. casei* DSM 20011 showed the greatest reduction in both MOSH and MOAH fractions. Relative to the control, this corresponded to a mean reduction of 77% for MOSH and 73% for MOAH. In comparison, *Lactiplantibacillus plantarum* 20174, *Lacticaseibacillus paracasei* DSM 20020, and *Lacticaseibacillus rhamnosus* DSM 20021 achieved mean MOSH reductions of 56%, 42%, and 64%, respectively, and MOAH reductions of 52%, 39%, and 63%, respectively. Based on these initial strain screening findings, *L. casei* DSM 20011 was selected for all subsequent experiments.

Although these initial strain screening results suggest strain-dependent differences in *in vitro* MOH removal efficiency, the screening was intended solely to identify the most promising candidate for further investigation. Additional studies, including a larger number of biological replicates and evaluating different MOH concentrations, bacterial cell densities, incubation times, and culture media, are required to confirm these observations. Such factors have previously been reported to influence the ability of probiotic bacteria to bind and remove chemical contaminants and toxins (Khorshidian *et al*., 2016; Serrano-Niño *et al*., 2014; Yousefi *et al.,* 2021; Zhao *et al*., 2013).

### 3.2 In Vitro removal of MOH by Lacticaseibacillus casei DSM 20011

Following the initial strain screening, the *in vitro* MOH removal assay was performed using *L. casei* DSM 20011 in eight independent biological replicates. A bacterial concentration of 10^8^ CFU/mL was selected based on the study of Yousefi *et al*. (2018), who investigated the ability of LAB to remove four PAH, namely benzo(*a*)pyrene (BaP), benz(*a*)anthracene (BaA), chrysene (Chr), and benzo(*b*)fluoranthene (BbF), from contaminated PBS. In that study, bacterial concentrations ranging from 10^7^ to 10^10^ CFU/mL were evaluated. Although PAH removal increased with increasing bacterial density, the improvement observed between 10^8^ and 10^10^ CFU/mL was relatively small, with a maximum increase on average of approximately 10%. Therefore, a concentration of 10^8^ CFU/mL was considered sufficient for the present study. The study by Yousefi et al. (2018) was selected as a relevant study because PAH are representative aromatic hydrocarbons related to MOAH and, like MOH, are highly hydrophobic compounds. It was therefore hypothesized that MOH removal by *L. casei* primarily occurs through adsorption to the bacterial cell surface rather than through metabolic degradation. This hypothesis is consistent with the current understanding of hydrocarbon biodegradation, which requires specialized enzymes such as alkane hydroxylases that have been described in hydrocarbon-degrading bacteria but are not recognized as part of the metabolic capacity of LAB (van Beilen and Funhoff, 2007; Ji et al., 2013). Previous studies further indicate that the removal of hydrophobic contaminants by lactic acid bacteria is predominantly mediated by adsorption to cell wall components rather than by enzymatic degradation (Zhao et al., 2013; Yousefi et al., 2018).

To evaluate the MOH removal capacity of *L. casei*, the GRAVEX mineral oil mixture was selected as a representative model contaminant because its chromatographic profile and chemical composition resemble those of mineral oil contamination typically associated with recycled paper and board used in food packaging.. GRAVEX is characterized by a hydrocarbon distribution centered around *n*-C18, extending approximately from *n*-C13 to *n*-C25, and contains about 25–27% MOAH. The final concentration used in the present study (0.0476 mg/mL) was selected to provide a food-relevant contamination level while ensuring reliable analytical quantification of MOH. In this context, Vollmer et al. (2011) analyzed 119 dry food products marketed in Germany and reported MOSH concentrations ranging from 3 to 31 mg/kg and MOAH concentrations from 0.5 to 6.1 mg kg ¹, with MOAH representing approximately 7–19% of the migrated mineral oil fraction. Although the experimental system employed in the present study does not replicate the complexity of food matrices, the selected GRAVEX composition and concentration provide a realistic model for investigating the interaction between *L. casei* and food-relevant MOH under controlled *in vitro* conditions.

The representative LC–GC-FID traces (**Figure 1**) visually confirm the quantitative results presented in Figures 2 and 3. Compared with the control, incubation with *L. casei* DSM 20011 resulted in a considerable reduction in both the MOSH and MOAH chromatographic profiles. Despite the reduced abundance, the overall shape and distribution of the unresolved hydrocarbon humps remained largely unchanged, indicating that *L. casei* DSM 20011 reduced both hydrocarbon fractions without causing major alterations to their chromatographic profiles. Therefore, no apparent preferential reduction of specific hydrocarbon fractions was observed, suggesting that the removal occurred uniformly across the analyzed MOSH and MOAH carbon number ranges. Nevertheless, these observations should be confirmed by comprehensive two-dimensional gas chromatography coupled with mass spectrometry (GC×GC–MS), which would enable a more detailed characterization of individual hydrocarbon classes and compounds.

**Figure 1:**
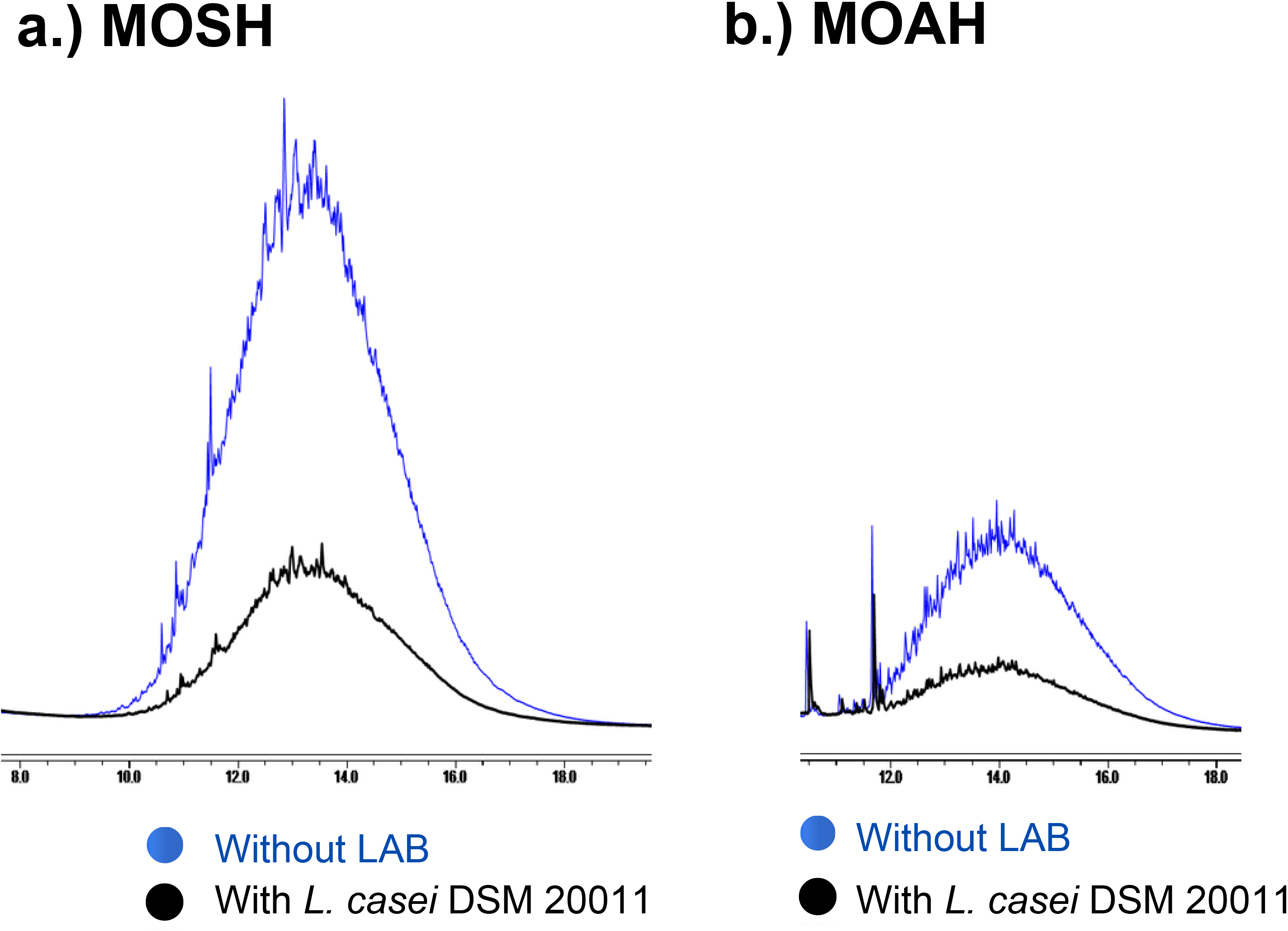
Representative LC–GC-FID traces of the a.) MOSH and b.) MOAH fractions following *in vitro* incubation in the absence (control: PBS supplemented with the MOH mixture, blue traces) and presence (PBS supplemented with the MOH mixture and *L. casei* DSM 20011, black traces) of *L. casei* DSM 20011.

**Figure 2:**
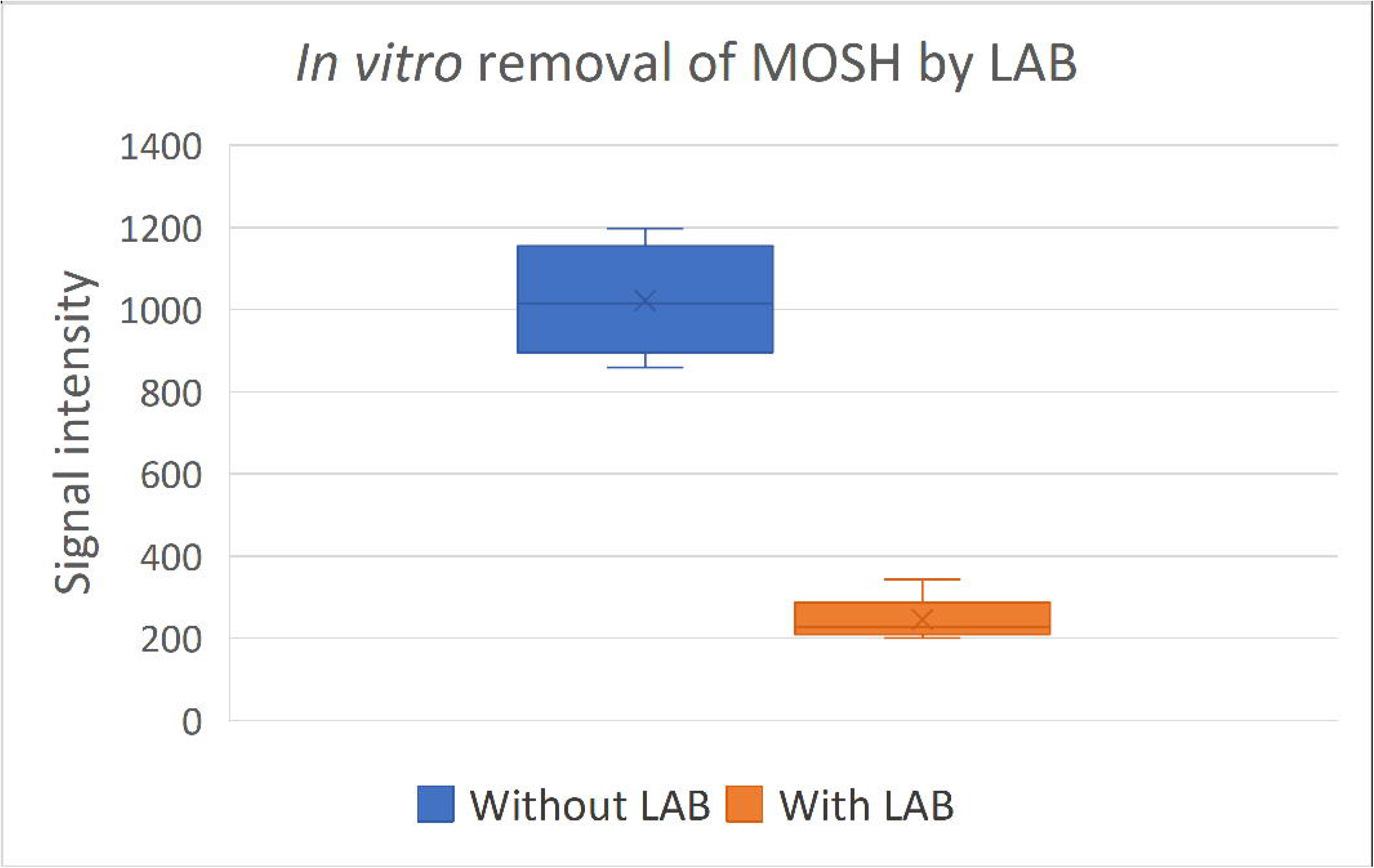
Box plot of *in vitro* removal of MOSH by lactic acid bacteria (LAB). **<u>Without LAB</u>**: Control sample – PBS with MOH mixture (n=4). **<u>With LAB</u>**: PBS with MOH mixture and *L. casei* DSM 20011 (n=8).

**Figure 3:**
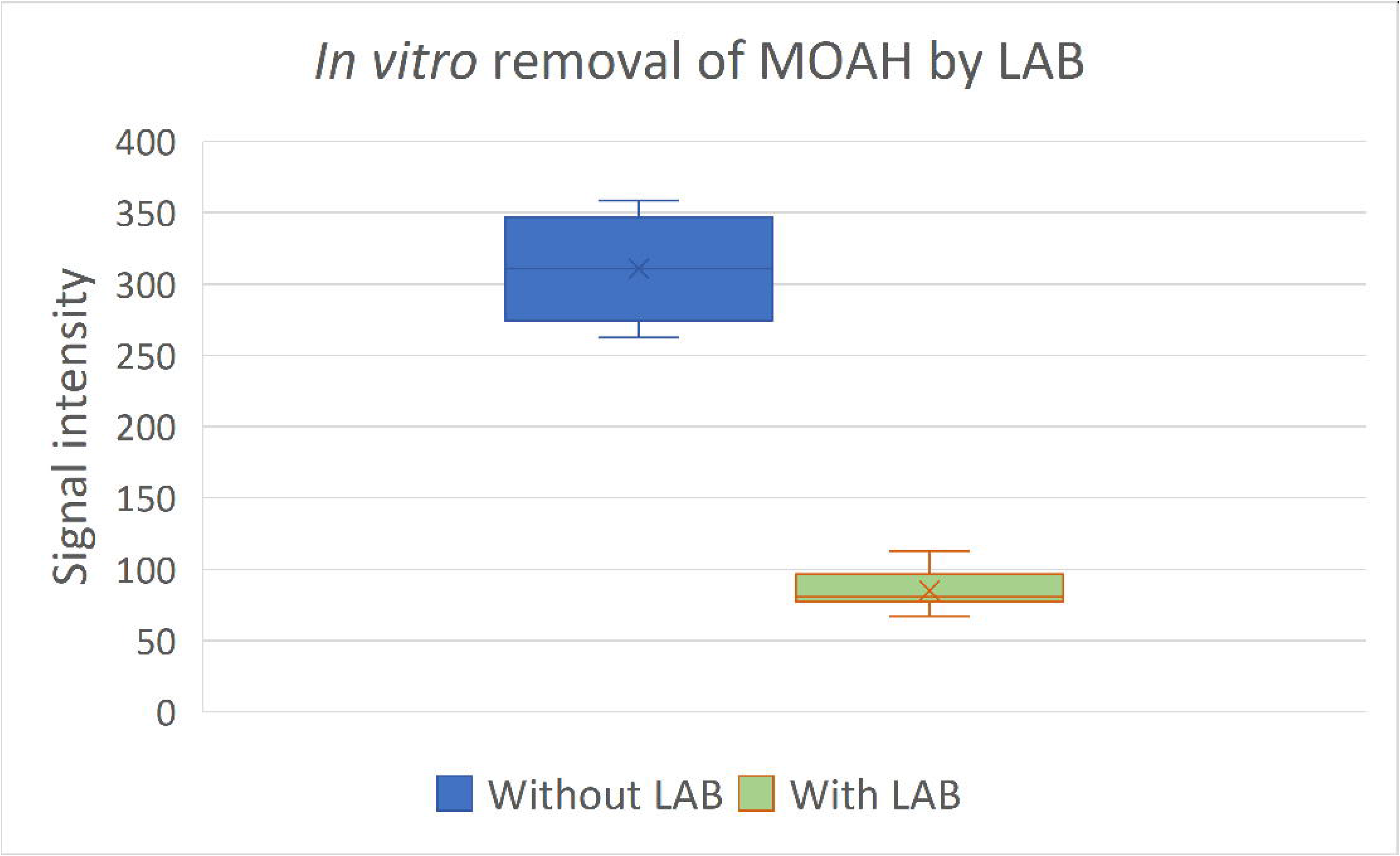
Box plot of *in vitro* removal of MOAH by lactic acid bacteria (LAB). **<u>Without LAB</u>**: Control sample – PBS with MOH mixture (n=4). **<u>With LAB</u>**: PBS with MOH mixture and *L. casei* DSM 20011 (n=8).

**Figures 2** and **3** show the signal intensity of MOSH and MOAH, respectively, determined by LC–GC-FID following *in vitro* incubation in the absence (control) or presence of *L. casei* DSM 20011. In both cases, incubation with *L. casei* resulted in a statistically significant reduction (*p* < 0.05) in MOH concentrations compared with the control. The concentrations of both MOSH and MOAH decreased by up to with a maximum observed reduction of approximately 80%, and mean reductions of 77 ± 5% and 73 ± 5%, respectively,demonstrating the high removal capacity of *L. casei* DSM 20011 under the experimental conditions employed. As no previous studies have investigated the removal of MOH by LAB, the present results were compared with the available literature on the binding of PAH. Zhao et al. (2013) evaluated the benzo[a]pyrene (BaP)-binding ability of 15 *Lactobacillus* strains and reported that *Lactiplantibacillus plantarum* CICC 22135 and *Lactiplantibacillus pentosus* CICC 23163 exhibited the highest binding capacities, reaching 66.8% and 64.3%, respectively. Similarly, Yousefi et al. (2018) reported the highest *in vitro* removal of PAH by *Lacticaseibacillus acidophilus* LA-5, with binding efficiencies ranging from 73% to 84% at a bacterial concentration of 10¹ CFU/mL. These findings indicate that the MOH removal capacity of *L. casei* DSM 20011 is within or exceeds the range of binding efficiencies previously reported for PAH by lactic acid bacteria.

Overall, the present study demonstrates that *L. casei* DSM 20011 efficiently removed both MOSH and MOAH *in vitro*, with no apparent preferential reduction of specific hydrocarbon fractions. As previously reported for PAH binding by LAB, contaminant removal appears to be strain-dependent (Yousefi *et al*., 2018), highlighting the need for further studies to elucidate the mechanisms underlying strain-specific differences and the apparently uniform removal of chemically diverse hydrocarbon fractions.

## 4. Conclusions

This study provides the first evidence that *L. casei* DSM 20011 can efficiently remove both MOSH and MOAH under *in vitro* conditions. The observed reduction of up to approximately 80% and the comparable removal across the analyzed hydrocarbon fractions highlight the potential of this food-derived bacterium as a promising candidate for reducing MOH contamination. The removal efficiency was within the upper range of binding capacities previously reported for PAH by LAB. Considering the structural relationship between PAH and the MOAH fraction of MOH, together with the absence of recognized hydrocarbon-degrading enzymes in LAB, adsorption to the bacterial cell wall appears to be the most plausible explanation for the observed MOH removal, although this hypothesis requires experimental confirmation. Further studies are required to experimentally confirm the proposed adsorption mechanism, investigate possible strain-specific differences in MOH removal, and evaluate the effectiveness of *L. casei* DSM 20011 in complex food matrices.

## Acknowledgments and funding sources

This work was supported by ByNewBioactive that was founded on the basis of a contract for the execution and (co)financing of a research project to “strengthen the international mobility of Slovenian researchers and research organizations and to promote the international involvement of Slovenian applicants” (action code C3.K8.IC) under the Recovery and Resilience Plan and based on evaluation summary report (ESR) under Horizon Europe International Call MSCA – HORIZON-MSCA-2022-PF-0.1. In addition, this work was supported by the European Union under Horizon Europe (project 101186975—ERA Chair FOODOMICS). The authors are grateful to Irina Pavlič for her assistance with the chemical analysis. The authors are grateful to Irina Pavlič for her assistance with the chemical analysis.

## Author contribution

AMV: Conceptualization, Investigation, Methodology, Project administration, Software, Validation, Visualization, Writing – original draft, Writing – review & editing, Resources, Funding acquisition. MP: Methodology. CC: Methodology. LB: Writing – review & editing. DP: Writing – review & editing. LI: Writing – review & editing, Supervision. SM: Writing – review & editing, Supervision.

## SUPPLEMENTARY MATERIAL

**Table S1:** Preliminary screening of different LAB strains for *in vitro* MOH removal. All strains and control samples were tested following the same erimental protocol.

| **LAB** | **MOSH**<br>(signal intensity\*) | **MOSH reduction\*\***<br>**(%)** | **MOAH**<br>(signal intensity) | **MOAH reduction\*\***<br>**(%)** |
|---|---|---|---|---|
| PBS with MOH mixture and ***Lactiplantibacillus plantarum*** **20174** | 450 ± 50 | 56 ± 5 | 150 ± 16 | 52 ± 5 |
| PBS with MOH mixture and ***Lacticaseibacillus paracasei*** **DSM 20020** | 596 ± 110 | 42 ± 10 | 190 ± 33 | 39 ± 10 |
| PBS with MOH mixture and ***Lacticaseibacillus rhamnosus*** **DSM 20021** | 364 ± 84 | 64 ± 8 | 116 ± 23 | 63 ± 7 |
| PBS with MOH mixture and ***Lacticaseibacillus casei*** **DSM 20011** | **237** ± 51 | **77 ± 5** | **84** ± 15 | **73 ± 5** |
| **Control 1**: PBS with MOH mixture without LAB | 1020 ± 140 | **-** | 310 ± 40 | **-** |
| **Control 2:** PBS without MOH mixture and LAB | 49 ± 12 | **-** | 14 ± 6 | **-** |

\*MOSH and MOAH are reported as mean chromatographic hump area signal intensities ± SD, because this screening

